# Stomatal density affects rice mesophyll cell size and shape and modulates a conserved pattern of cells through the leaf

**DOI:** 10.1101/2022.11.09.515764

**Authors:** Sloan Jen, Im-Chai Saranrat, Qi Yang Ngai, Xiao Yi, Armand Jodie, Matthew J. Wilson, Zhu Xin-Guang, Andrew J. Fleming

**Affiliations:** Plants, Photosynthesis and Soil, School of Biosciences, University of Sheffield, Sheffield, UK; Carl R. Woese Institute for Genomic Biology, University of Illinois at Urbana Champaign, Urbana, IL, USA; Institute of Plant Physiology & Ecology, Shanghai Institutes for Biological Sciences, Chinese Academy of Sciences, Shanghai, China

**Keywords:** Cell shape, cell size, CO_2_, light, mesophyll, photosynthesis, rice, stomatal density, stomatal conductance

## Abstract

The structure of the mesophyll influences how light, CO_2_ and water travels inside a leaf, affecting the rates of both photosynthesis and transpiration. Recent studies in wheat and Arabidopsis have shown that the structure of the mesophyll is influenced by the density and activity of stomata, consistent with the hypothesis that gas flow via stomata can modulate internal cell growth and separation to co-ordinate leaf structure and function. To investigate whether this also occurs in rice, a staple food crop for a large fraction of the world’s population, we examined mesophyll structure in rice mutants with altered stomatal density. Our data show that stomatal function modulates mesophyll structure in rice. Variation in the degree of mesophyll lobing made a major contribution to altered mesophyll structure, suggesting that modified leaf gas flux through stomata influences an aspect of cell shape directly linked to gas exchange capacity in rice. In addition, our analysis revealed a previously unreported underlying pattern in cell size, shape and axiality across layers of the rice mesophyll, which further investigation revealed is present in a range of rice species and cultivars. The potential origin and significance of this mesophyll patterning are discussed.

**HIGHLIGHT:** We describe a previously unreported cellular pattern in rice leaves and show that it is modulated by stomata. These results shed new light on leaf structure and function.

## INTRODUCTION

Sandwiched between the upper and lower surfaces of the leaf is the mesophyll – the main site of photosynthesis. In dicotyledonous plants the mesophyll is typically separated into two layers: the adaxial tall, vertically orientated palisade cells, and the abaxial less organised, irregularly shaped ‘spongy’ mesophyll cells (Esau, 1965; Pyke, 2012). However, in monocotyledonous plants the mesophyll has traditionally been seen as more uniform (Chonan, 1978). The photosynthetic capacity of a leaf is intrinsically linked to the structure of its mesophyll. To reach a chloroplast for photosynthetic fixation, atmospheric CO_2_ must diffuse into the leaf via the stomata, through the intercellular airspace and into the mesophyll cells across their cell walls. The area of mesophyll cell wall exposed to intercellular air space (*S*_*mes*_) and the relative proportions of air, cell, and cell wall within the mesophyll can determine its resistance to CO_2_ diffusion (Evans, 2021), which limits photosynthesis (Flexas *et al*., 2012). In a similar way, mesophyll structure influences the rate of water loss during transpiration, as CO_2_ and water travel in opposite directions along a common pathway through the mesophyll (Wong *et al*., 2022). Light movement through the leaf is also affected by the shape of mesophyll cells, with elongated palisade cells facilitating the penetration of light deeper into the leaf, and the more irregularly shaped spongy mesophyll cells helping to scatter light and maximise absorption (Vogelmann and Evans, 2002; Johnson *et al*., 2005).

To maximise photosynthetic capacity the internal structure of a leaf must be coordinated with its external environment during development. Regulation of stomatal density and size in response to the environment is well understood (Casson and Gray, 2008), and structure of the mesophyll has also been shown to respond to the conditions under which the leaf develops. For instance, higher temperatures can lead to a thinner mesophyll tissue (Habermann *et al*., 2022), and low light drives reduced cell expansion and division (with one fewer layer of cells in the palisade mesophyll), leading to a thinner mesophyll than in leaves grown under high light (Kalve *et al*., 2014; Hoshino *et al*., 2019).

Recent work suggests that stomatal function may influence mesophyll differentiation, with a potential link between leaf gas flux and mesophyll surface area. Dow, Berry and Bergmann (2017) showed that mesophyll cell density positively correlates with stomatal density in Arabidopsis *epf* mutants. Furthermore, Lundgren et al., (2019) showed that the porosity of the palisade mesophyll is higher in transgenic Arabidopsis plants with increased stomatal density and correlates positively with stomatal conductance (*g*_*s*_). This phenomenon is not exclusive to eudicots, with the same study determining that the correlation between stomatal density and *g*_*s*_ remains across a range of wheat species.

The experiments described above linking stomatal function to mesophyll structure have been performed on both monocot grasses (wheat) and eudicot (Arabidopsis) suggesting it may be a conserved mechanism to coordinate leaf structure and function. However, this is yet to be reported in other important crops, such as rice. The classic histology of the rice leaf is well established, with numerous papers describing the basic cell types (for example, mesophyll, bundle sheath, epidermal, stomata, xylem, phloem) and how these cell types are arranged in space to form the tissues that constitute the leaf (Esau, 1965; Chonan, 1978). Several studies have also considered how the arrangement of different cell and tissue types might contribute to overall leaf function, particularly in terms of photosynthesis; for example the efficiency of light capture, gas flux, and transport of the products of photosynthesis (Vogelmann, 1993; Parkhurst, 1994; Xiao and Zhu, 2017). The rice mesophyll is of particular interest because of the special role that mesophyll cell lobing may play in increasing the cell surface area available for photosynthetic gas exchange (Sage and Sage, 2009).

In this paper, we report on experiments that investigate the influence of altered stomatal density on mesophyll structure in rice. Our results suggest that stomatal function modulates mesophyll structure in rice mostly via altered mesophyll cell lobing, a parameter which, via its relationship to cell surface area to volume ratio, is expected to influence leaf gas exchange capacity. Interestingly, our analysis also revealed a previously unreported pattern in cell size, shape and axiality across layers within the rice mesophyll - the potential significance of this patterning is discussed.

## MATERIALS AND METHODS

### Plant material and growth conditions

OsEPF1OE-W, OsEPF1OE-S, OsEPFL9OE-2 and OsEPFL9OE-3 and their *Oryza sativa* (IR64) controls were kindly gifted to us by Professor Julie Gray. *O. sativa* (Indica) MR220, *O. sativa* (fragrant) MRQ76, and *O. sativa* (Indica) Malinja were provided by the Malaysian Agricultural Research and Development Institute. *Oryza punctata, Oryza meridionalis* and *Oryza latifolia* were provided by the International Rice Research institute. Rice plants were grown in a Conviron controlled environment chamber at 70% relative humidity, in a 12hr/12hr light/dark cycle at 28°C/24°C with a light intensity of 750 μmol m^−2^ s^−1^ at canopy height. Plants were germinated on filter paper with 15 ml water in petri dishes, then grown in 13D pots (0.88L) filled with 71% Kettering Loam (Boughton, UK), 23.5% Vitax John Innes No. 3 (Leicester, UK), 5% silica sand and 0.5% Osmocote Extract Standard 5–6 month slow-release fertilizer (ICL, Ipswich, UK) by volume, saturated with water for 4 to 5 weeks before gas exchange analysis was carried out and leaf samples were collected for imaging.

### Stomatal conductance

Stomatal conductance was measured using a LI-600 porometer (LI-COR, Lincoln, USA) set to a flow rate of 150 μmol s^-1^. Measurements were taken 2-3 hours into the light period, on the middle portion of fully expanded leaf 6, 28 days after sowing. Abaxial and adaxial conductance was measured and a mean taken of the two values.

### Microscopy

All samples were taken from the middle 3cm portion of the fully expanded leaf 6. OsEPF1-OE plants were harvested 21 days after sowing. All other plants were harvested 28 days after sowing. Abaxial epidermal stomatal densities and sizes were measured on nail varnish peels of dental resin impressions, with 4 fields of view per replicate. Images were taken on an Olympus BX51 microscope with an Olympus DP71 camera. Measurement of guard cell length and calculation of *g*_*smax*_ was performed as in (Caine *et al*., 2019) on 20 stomata per plant (five per field of view) from six biological replicates.

Samples for Technovit® sectioning (Fig. 1 and 2) and fresh transverse hand sections (Fig. 3 and 4) were fixed in 1:4 acetic anhydride:ethanol for 48 hours, then transferred to 70% ethanol. Hand sections were cleared in chloral hydrate saturated lactic acid for 2 hours at 70°C, then stained for 20-30 seconds with 0.05% Toluidine Blue O. Technovit® samples were embedded in Technovit® 3040 resin and sectioned at 7μm using a Leica Microtome, then stained for 20 seconds with Toluidine Blue O. All images were observed using an Olympus BX51 light microscope, with the 40x objective, Olympus DP71 camera and Cell B imaging software. Regions of interest were between the first and second major vein out from the mid vein, between two minor veins.

**Figure 1:**
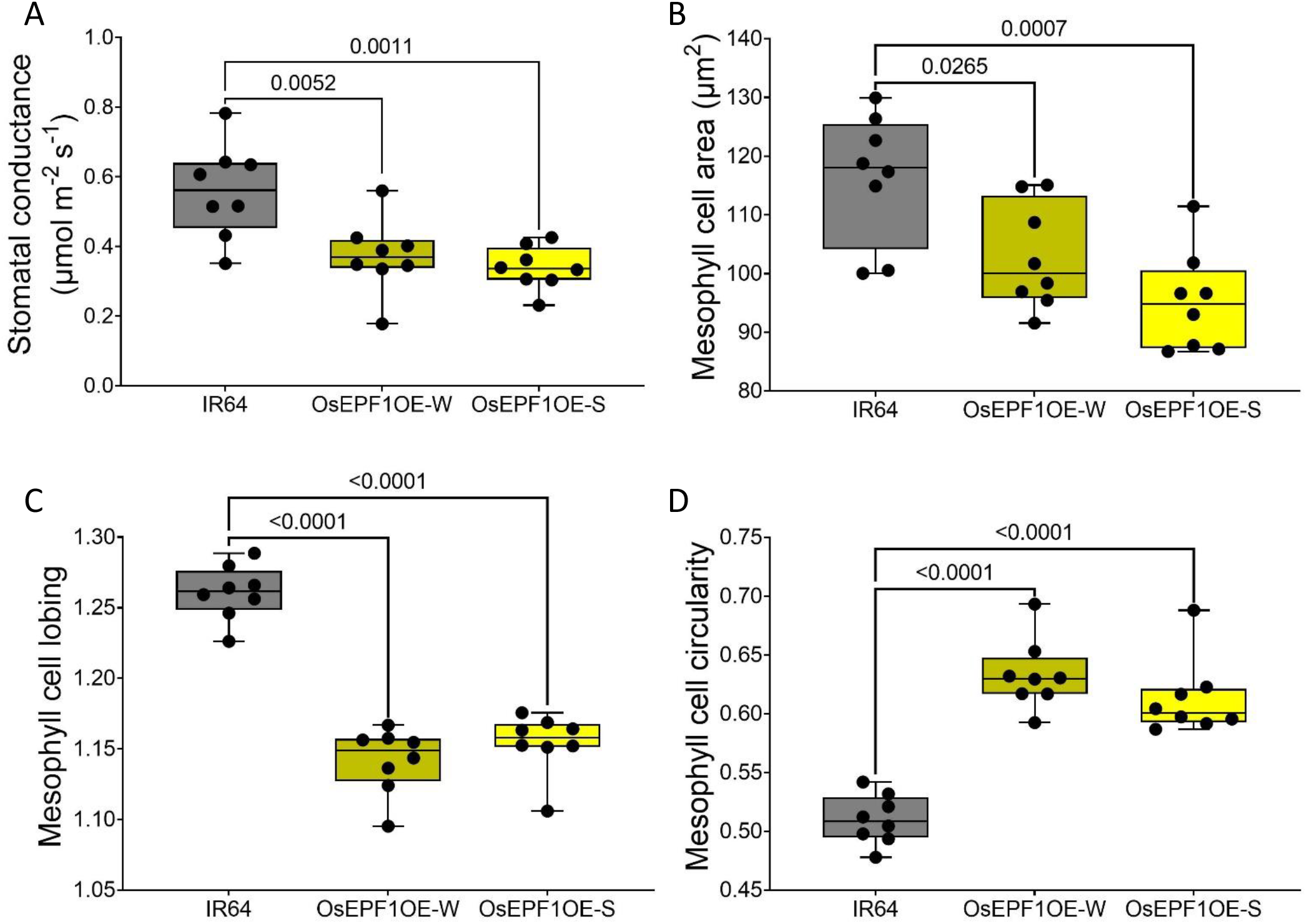
Reducing stomatal conductance affects mesophyll cell size and shape. Data from the middle of leaf 6 of 28 day old plants. OsEPF1OE weak (W) and strong (S) lines, and their IR64 control. **A)** OsEPF1OE stomatal conductance is significantly lower than in the control. One way ANOVA, p < 0.0001, n = 8. **B)** Mesophyll cell area is significantly lower in OsEPF1OE lines. One way ANOVA, p < 0.0001, n = 8. **C)** Mesophyll cell lobing is significantly lower in OsEPF1OE. One way ANOVA, p < 0.0001, n = 8. **D)** Mesophyll cell circularity is significantly higher in OsEPF1OE lines. One way ANOVA, p < 0.0001, n = 8. All multiple pairwise comparisons, Tukey, p values as shown, n = 8.

**Figure 2:**
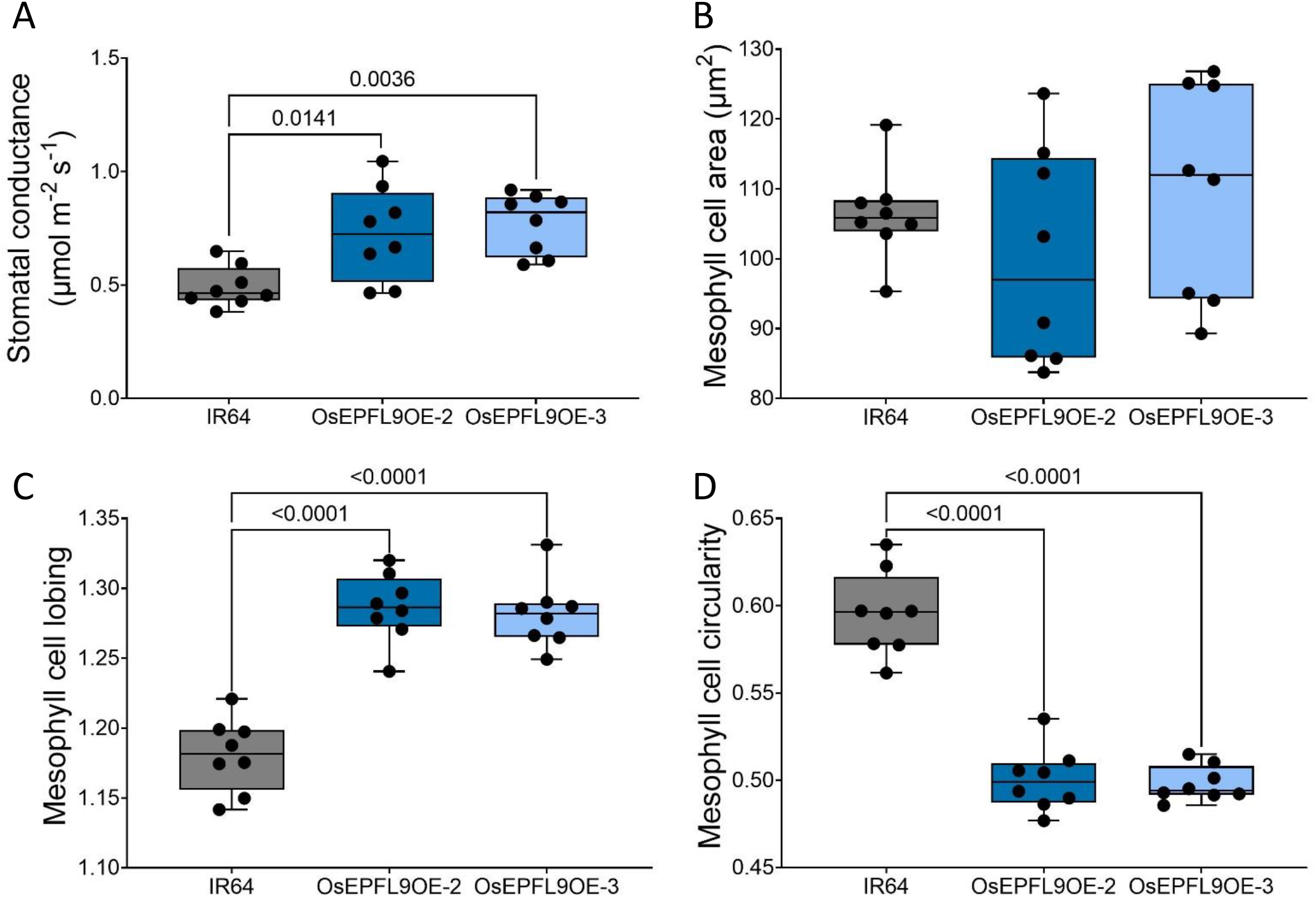
Mesophyll cell shape is affected by increased stomatal conductance. Data from the middle of leaf 6 of 21 day old plants. Two individual EPFL9OE lines (2 and 3) and their IR64 control. **A)** OsEPFL9OE stomatal conductance is significantly higher than the control line. one way ANOVA, p = 0.0028, n = 8. **B)** Mesophyll cell area is not affected by the change in stomatal conductance. One way ANOVA, p = 0.3389, n = 8. **C)** Mesophyll cell lobiness is significantly higher in both OsEPFL9OE lines. One way ANOVA,, p < 0.0001, n = 8. **D)** Mesophyll cell circularity is significantly lower in OsEPFL9OE plants. One way ANOVA, p < 0.0001, n = 8. All multiple pairwise comparisons, Tukey, p values as shown, n = 8.

**Figure 3:**
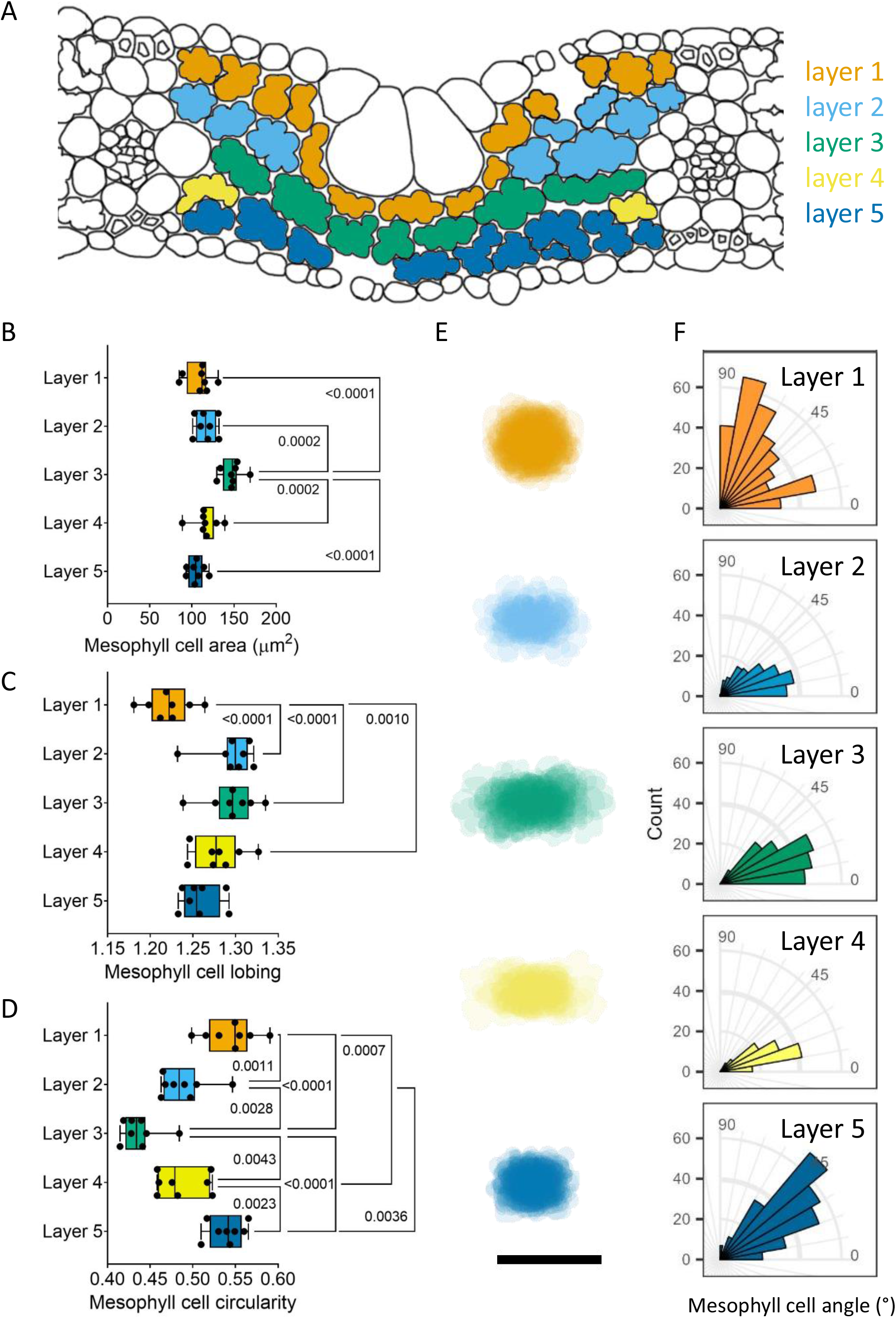
The rice mesophyll can be divided into 5 cell tissue layers. **A)** Representation of rice mesophyll with different cell layers highlighted from layer 1 (touching the adaxial epidermis) to layer 5 (touching the abaxial epidermis). Layer 3 is a continuous row of cells between the two minor veins. **B-F)** Representative data from middle of leaf 6 of 28 day old IR64 control (from EPF1OE experiment). **B)** Mesophyll cell area is largest in layer 3 **C)** Mesophyll cell lobing is lowest in layer 1. **D)** Mesophyll cell circularity is lowest in layer 3. **B-D)** One way ANOVA p < 0.0001, Tukey’s multiple comparison test, p values as shown, n = 8. **E)** Mesophyll cell projections of all cells in each layer from one representative individual. **F)** Mesophyll cell angle – the angle of the longest axis of each cell differs by cell layer. Scale bar = 20 μm

**Figure 4:**
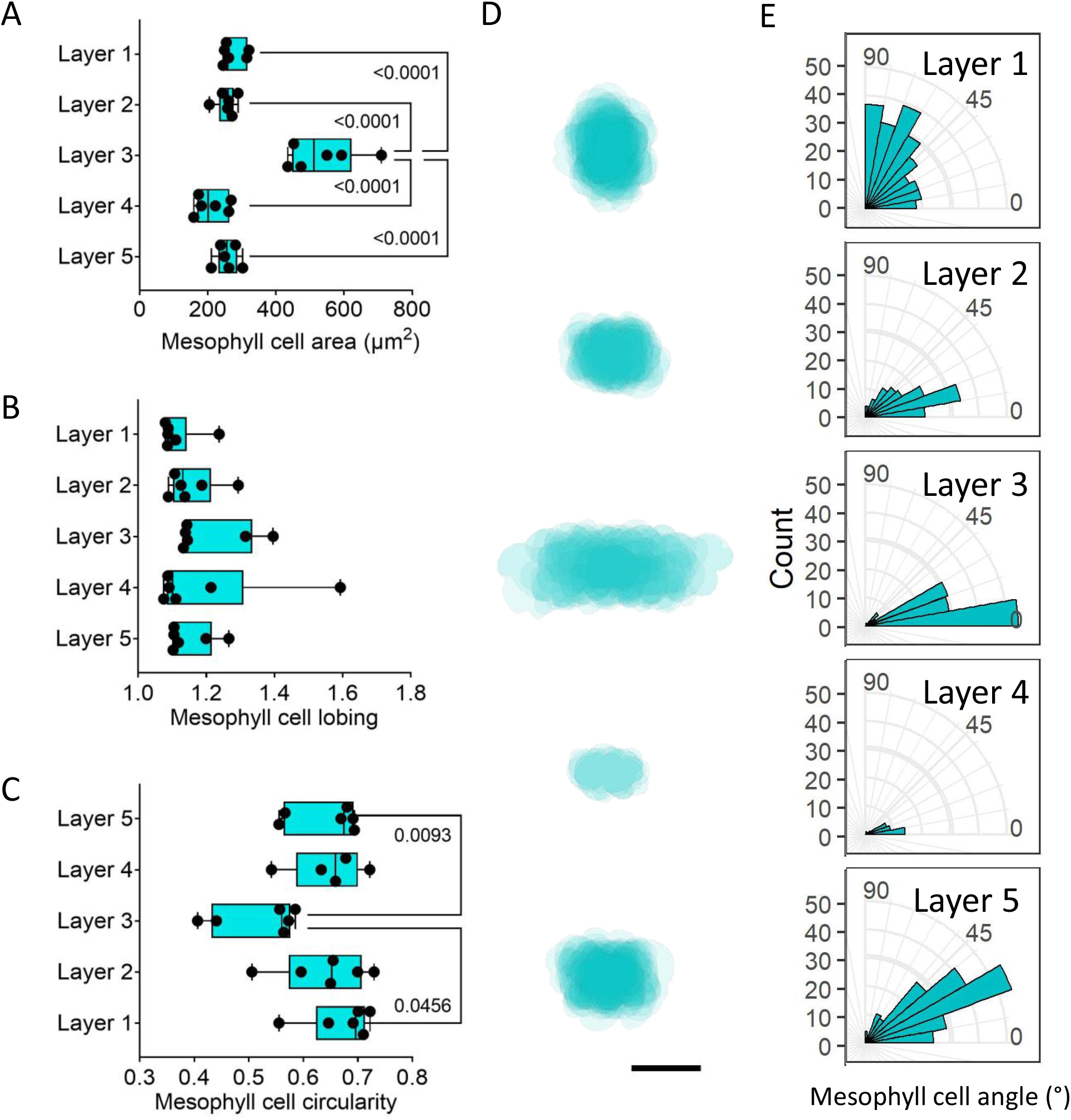
The tissue layer patterning seen in IR64 is present in a range of rice varieties – demonstrated by *O. latifolia*. Representative data from middle of leaf 6 of 28 day old *O. latifolia*. **A)** Mesophyll cell area is largest in layer 3, One way ANOVA, p < 0.0001, Tukey pairwise multiple comparisons, p values as shown, n = 6 **B)** Mesophyll cell lobing is lowest in layer 1. **C)** Mesophyll cell circularity is lowest in layer 3. One way ANOVA, p = 0.0105, Tukey pairwise multiple comparisons, p values as shown, n =6. **D)** Mesophyll cell projections of all cells in each layer from one representative individual. **E)** Mesophyll cell angle – the angle of the longest axis of each cell differs by cell layer. Scale bar = 20 μm

Mesophyll cell image analysis was performed in FIJI (ImageJ 5.3g) software using an in-house macro. The mesophyll layers were identified relative to their position in the leaf (**Fig. 3A**). Layer 1 was identified as directly below the upper epidermis and bulliform cells, layer 3 linking the middle of the left and right minor vein, layer 5 directly above the lower epidermis, layer 2 between layers 1 and 3, and layer 4 between layers 3 and 5. Every cell within the layer was outlined by hand, and area (μm^2^), perimeter (μm), circularity, cell length (Feret), cell width (MinFeret), convex hull perimeter (μm) and cell angle (FeretAngle) measurements taken. Mesophyll cell lobing was calculated as cell perimeter divided by convex hull perimeter, FeretAngle measurements were adjusted so that 0° is in line with a line between the minor veins in the image, and 90° is perpendicular to that line (see **Supplementary Fig. S7** at *JXB* online for details). Cell projection images were created using an in-house FIJI macro – first the long axis of each cell was rotated to horizontal, then cell outlines were superimposed. For OsEPF1OE and OsEPFL9OE lines, leaf sections from eight plants were imaged. For each rice species/variety in **Fig. 3** and **4**, leaf sections from four to six different plants were imaged. From each biological repeat, four images were analysed.

### Computational modelling

To explore the potential impacts of larger mesophyll cells in the middle layer to leaf photosynthesis, we built four simplified models of mesophyll cell packing (**Fig. 5A-D**). Model 1 adopted a mix of two cell types with larger cells in its middle layer. Cell length and cell width was rounded based on measurements from O. latifolia, so that total length of three large cells in a layer equals to the total length of five small cells, and total thickness of four large cells equals to the total thickness of five small cells (**Supplementary Fig. S15**). In this way, Model 2 was generated by replacing the middle layer in model 1 with small cells, and Model 3 has a same leaf thickness as Model 2. Thickness of plastid layer in both cell types were calculated by keeping the same plastid volume between a layer of larger cells and a layer of small cells. Model 4, therefore, has the same plastid volume as Model 2 and Model 1. Size of vacuole in both cell types were also adjusted to result the same cytosol volume in Model 1, 2 and 4 (**Fig. 5E**).

**Figure 5:**
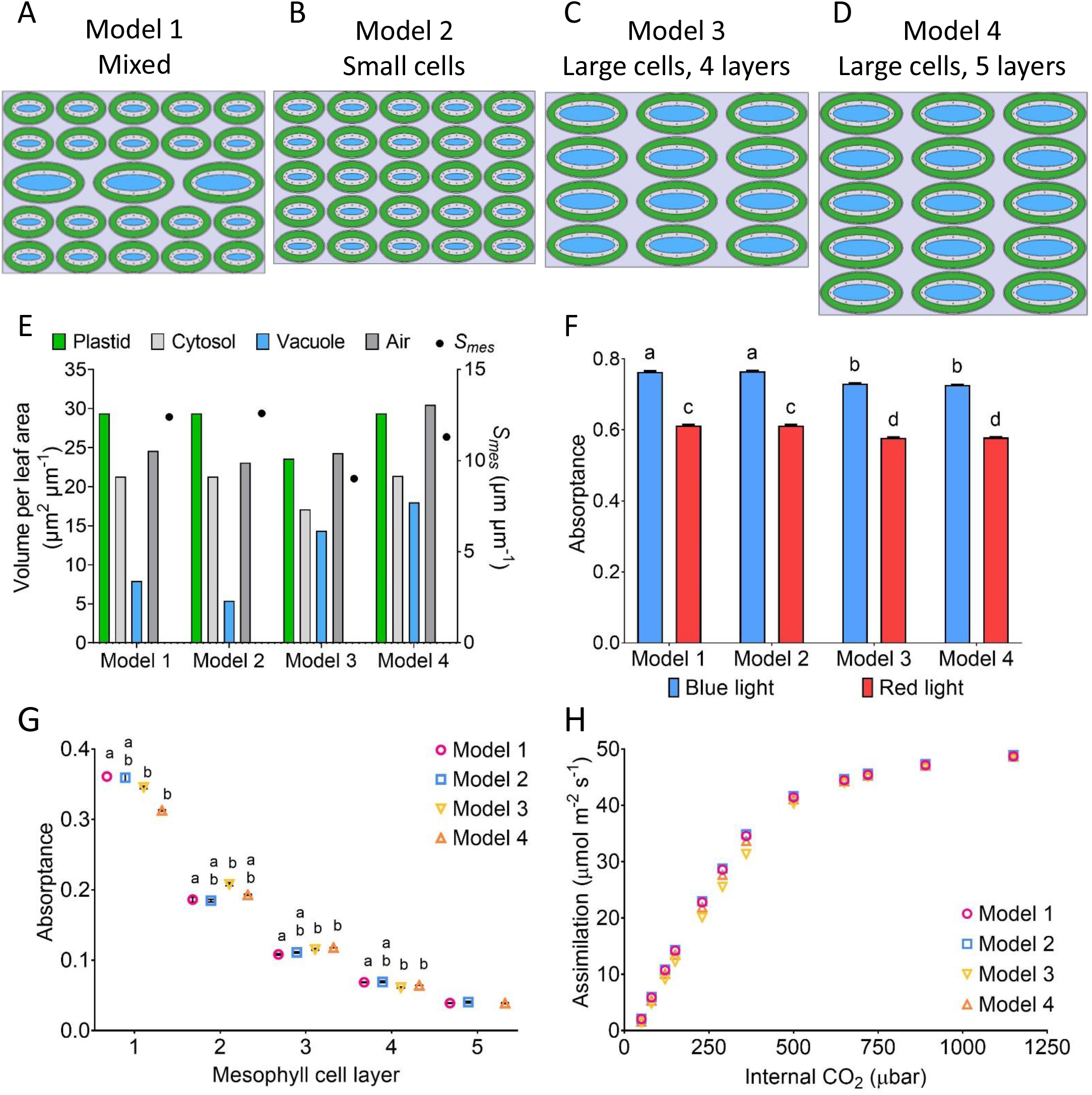
CO_2_ and light move differently through four simplified mesophyll tissue models. Four cell tissue layer models were designed, green represents plastid, white centres represent cytosol: **A)** Model 1 has larger cells in the middle layer (layer 3), **B)** Model 2 has five layers of small cells, **C)** Model 3 has four layers of large cells, **D)** Model 5 has five layers of large cells. Models 1, 2 and 4 have the same plastid and cytosol volume. Models 2 and 3 are the same leaf thickness. **E)** *S*_*mes*_ and the proportions of different cell elements in the 4 models. F) Total red and blue light absorptance is higher in Models 1 and 2 than Models 3 and 4 – mean with SEM. Two way ANOVA, p< 0.0001, Tukey multiple comparison - different letters represent significantly different values, p < 0.0001, n = 3 **G)** Blue light absorptance in each cell layer of the 4 models – mean values with SEM. Individual one way ANOVA performed for each cell layer - Layers 1-4 p < 0.001, Layer 5 ns, Tukey multiple comparison - different letters represent significantly different values, p < 0.05, n = 3 H) Assimilation/Internal CO_2_ (*C*_*i*_) curves are very similar for the four models. Mean values, n = 3, SEM is too small for error bars to show.

With the constructed leaf geometry, light propagation inside the leaf was simulated by a Monte-Carlo ray tracing algorithm (Govaerts *et al*., 1996; Xiao *et al*., 2016, 2022). Due to the neglect of epidermis cells here for these simplified models, diffuse incident rays were emitted onto the upper boundary as the light source. Density of rays were tested to ensure the convergence of simulation, which is also reflected by the small error bars in the predicted light absorptance by the whole leaf (**Fig. 5F**) and each layer (**Fig. 5G**). Light absorptance of each chloroplast under blue and red light were simulated and applied to the later calculation of carboxylation rate for the process of CO_2_ reaction and diffusion. Details of the ray tracing algorithm and a list of related parameters can be found in the Supplementary Material.

Process of CO_2_ reaction and diffusion inside the leaf was simulated by a partial differential system (Tholen and Zhu, 2011; Xiao and Zhu, 2017; Xiao *et al*., 2022). A constant CO_2_ concentration ([CO_2_]) was set to the upper and lower boundaries, representing [CO_2_] in the substomatal cavity, i.e. C_i_. Inside the compartments of air space, cytosol, chloroplast, mitochondria and vacuole, reaction-diffusion processes of CO_2_ were modeled by the following equations:

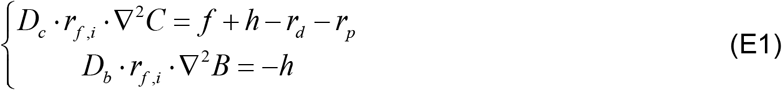

where *C* (mol m^-3^) and *B* (mol m^-3^) are the concentration of CO_2_ and HCO_3_^-^ respectively. *D*_*c*_ (m^2^ s^-1^) and *D*_*b*_ (m^2^ s^-1^) are the liquid-phase diffusion coefficient of CO_2_ and HCO_3_^-^ in water correspondingly. *r*_*f,i*_ is a dimensionless factor representing the change of the diffusion coefficient relative to free diffusion in water in different compartments. ▽^2^*C* is the Laplace operator which equals 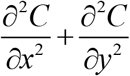. While on the right-hand side of the equation, *f* is volumetric carboxylation rate (mol m^-3^ s^-1^), *h* is hydration rate from CO_2_ to HCO_3_^-^ catalyzed by CA, and *r*_*d*_ is volumetric respiration rate, and *r*_*p*_ is volumetric photo-respiration rate. In addition, these terms are distributed differently in each compartment, for example, in the cytosol *f* = *r*_*d*_ = *r*_*p*_ = 0, in the chloroplast *r*_*d*_ = *r*_*p*_ = 0, and in mitochondria *f* = 0. The volumetric carboxylation rate and photo-respiration rate were calculated based on the Farquhar-von Caemmerer-Berry model (Von Caemmerer, 2013). Details of the reaction-diffusion system and parameters used can be found in the Supplementary Material.

## RESULTS

### Changing stomatal density leads to altered mesophyll cell size and shape

To investigate whether a relationship exists between stomatal function and mesophyll structure in rice, we exploited the availability of published rice transgenics with altered stomatal density. In OsEPF1OE plants an epidermal patterning factor (EPF1) has been overexpressed, leading to decreased stomatal density and a concomitant decrease in stomatal conductance (Caine *et al*., 2019). Two lines were investigated, OsEPF1OE-W, which has been reported to have a weak phenotype, and OsEPF1OE-S with a strong phenotype. Both OsEPF1OE-W and OsEPF1OE-S lines have a significantly lower stomatal conductance than the comparable IR64 control plants (**Fig. 1A**, one way ANOVA, p = 0.0008, Tukey multiple comparison test, p < 0.05, n = 8). Stomatal density is significantly reduced in OsEPF1OE lines, while stomata size is not altered, resulting in significantly decreased theoretical maximum stomatal conductance (*g*_*smax*_) (**Supplementary Fig. S1A**,**C**,**E**). To investigate mesophyll structure in these lines, we first measured mesophyll cell area. As shown in **Fig. 1B**, OsEPF1OE-W and OsEPF1OE-S plants have significantly smaller mesophyll cells (**Fig. 1B**, one way ANOVA, p < 0.0001, Tukey multiple comparison test, p = 0.0265, p = 0.0007, n = 8). We used two parameters to measure cell shape: circularity - which describes the similarity of a given shape to a circle, with a higher value denoting a rounder shape, and cell lobing – calculated from the perimeter of the cell, where a greater value represents a larger deviation from the perimeter of the convex hull (**Supplementary Fig. S12 B**). Mesophyll cell shape was also altered, with decreased cell lobing (**Fig. 1C**, one way ANOVA, p < 0.0001, Tukey multiple comparison test, p < 0.0001, n = 8) and increased cell circularity (**Fig. 1D**, one way ANOVA, p < 0.0001, Tukey multiple comparison test, p < 0.0001, n = 8) in both the OsEPF1OE-W and OsEPF1OE-S mesophyll.

To see if leaves with increased stomatal conductance also displayed a mesophyll phenotype, two transgenic rice lines overexpressing EPFL9 were used: OsEPFL9OE-2 and OsEPFL9OE-3 (Bertolino *et al*., 2022). Compared to IR64, these lines had a significantly higher stomatal conductance (**Fig. 2A**, one way ANOVA, p = 0.0028, n = 8, Tukey multiple comparison test, p < 0.05) and stomatal density (**Fig S1B**) but a slightly smaller stomata size (**Fig S1D**), resulting in a significantly increased *g*_*smax*_ (**Fig. S1F**). Mesophyll cell area was not significantly different in the transgenic leaves from the control IR64 plants (**Fig. 2B**, one way ANOVA, p = 0.3389, n = 8). However, there was a change in mesophyll cell shape with OsEPFL9OE plants having significantly increased lobing (**Fig. 2C**, one way ANOVA, p < 0.0001, Tukey multiple comparison test, p < 0.0001, n =8) and decreased cell circularity (**Fig. 2D**, one way ANOVA, p < 0.0001, Tukey multiple comparison test, p < 0.0001, n = 8).

### Analysis of WT and EPF mutants reveals linkage of cell size and shape to leaf layer position

During our analysis of the rice mesophyll, it became apparent that there was a non-random distribution of mesophyll cell size. In particular, it appeared that there might be a relationship between cell size and leaf layer position. To test this hypothesis, we assigned cells to tissue layers (1-5) as shown in **Fig. 3A**. To investigate this pattern, the cell size and shape data from the rice lines reported in **Fig. 1** and **Fig. 2** (IR64, OsEPF1OE, OsEPFL9OE) was split into the five tissue layers and analysed separately. For simplicity, data from the EPF1 IR64 control plants is shown in **Fig. 3** and the other five lines are shown in the supplementary material as the same pattern is seen in all six plant lines (**Supplementary Fig. S2-4**). Cell area varied significantly by layer (**Fig. 3B, Supplementary Fig. S2, Table 1**, one way ANOVA, p < 0.0001 or p = 0.0001, n = 8), with the cells in the middle layer (layer 3) being significantly larger than cells in all other layers (**Table 1**, Tukey multiple comparison test, p < 0.05, n = 8). This pattern was present in all the lines analysed, suggesting that the changes in cell size reported in **Fig. 1** and **Fig. 2** were superimposed on an endogenous pattern, which is maintained in rice lines with altered stomatal conductance.

**Table 1:**
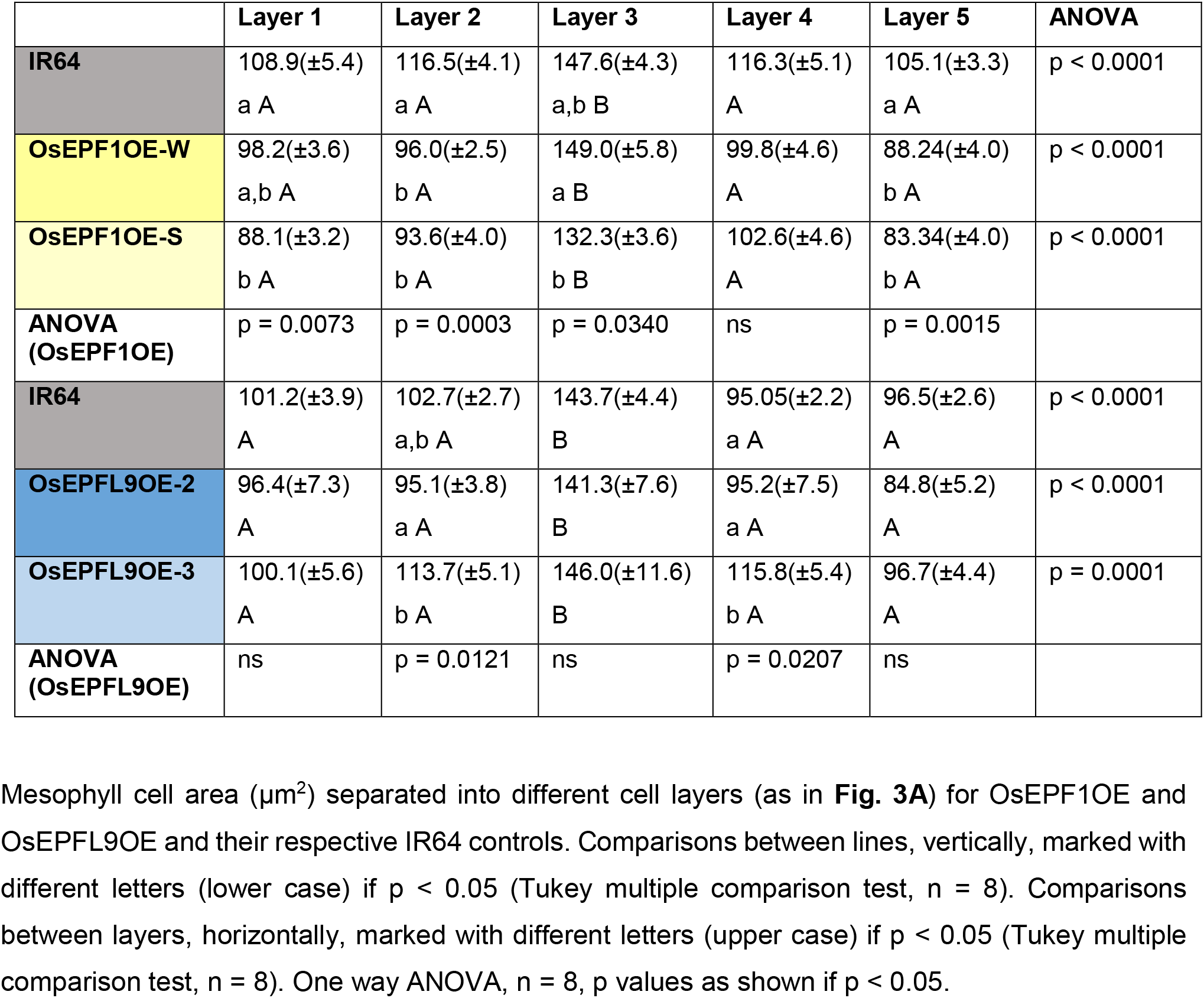
OsEPF1OE and OsEPFL9OE mesophyll cell area varies by cell layer, with layer 3 consistently being the largest. The pattern of cell area for each cell layer between plant lines follows the pattern shown in the total mesophyll.

|  | Layer 1 | Layer 2 | Layer 3 | Layer 4 | Layer 5 | ANOVA |
| --- | --- | --- | --- | --- | --- | --- |
| IR64 | 108.9(±5.4)<br>a A | 116.5(±4.1)<br>a A | 147.6(±4.3)<br>a,b B | 116.3(±5.1)<br>A | 105.1(±3.3)<br>a A | p < 0.0001 |
| OsEPF1OE-W | 98.2(±3.6)<br>a,b A | 96.0(±2.5)<br>b A | 149.0(±5.8)<br>a B | 99.8(±4.6)<br>A | 88.24(±4.0)<br>b A | p < 0.0001 |
| OsEPF1OE-S | 88.1(±3.2)<br>b A | 93.6(±4.0)<br>b A | 132.3(±3.6)<br>b B | 102.6(±4.6)<br>A | 83.34(±4.0)<br>b A | p < 0.0001 |
| ANOVA<br>(OsEPF1OE) | p = 0.0073 | p = 0.0003 | p = 0.0340 | ns | p = 0.0015 |  |
| IR64 | 101.2(±3.9)<br>A | 102.7(±2.7)<br>a,b A | 143.7(±4.4)<br>B | 95.05(±2.2)<br>a A | 96.5(±2.6)<br>A | p < 0.0001 |
| OsEPFL9OE-2 | 96.4(±7.3)<br>A | 95.1(±3.8)<br>a A | 141.3(±7.6)<br>B | 95.2(±7.5)<br>a A | 84.8(±5.2)<br>A | p < 0.0001 |
| OsEPFL9OE-3 | 100.1(±5.6)<br>A | 113.7(±5.1)<br>b A | 146.0(±11.6)<br>B | 115.8(±5.4)<br>b A | 96.7(±4.4)<br>A | p = 0.0001 |
| ANOVA<br>(OsEPFL9OE) | ns | p = 0.0121 | ns | p = 0.0207 | ns |  |

With respect to cell shape, analysis of cell lobing also identified variation between the leaf layers (**Fig. 3C, Supplementary Fig. S3, Table 2**, one way ANOVA, p = 0.014 – p < 0.0001, n = 8), although in this case layer 3 was not the most distinctive. Rather, cells in layer 1 (adaxial layer) of the mesophyll were generally distinguishable as having the lowest lobing value, with the lobing values in the other four layers generally being similar to each other. When mesophyll cell shape was calculated based on circularity (**Fig. 3D, Supplementary Fig. S4, Table 3**), a clear pattern emerged in which cells in the middle layer (layer 3) were significantly less circular than cells in the other layers - this was true in all six lines analysed (one way ANOVA, p < 0.0001, n = 8, Tukey multiple comparison test, p < 0.05 – p < 0.0001, n = 8). General differences in mesophyll cell size and shape between layers can be seen by projecting the cell shapes within each layer on top of each other (**Fig. 3E, Supplementary Fig. S5**). Cells in layer 3 appear less circular (more ellipsoid) and larger. Layer 1 cells are circular, while layer 5 cells are smaller and squarer.

**Table 2:** *OsEPF1OE* and *OsEPFL9OE* mesophyll cell lobing varies by cell layer, with layer 1 consistently having the lowest lobing. Cell lobing is consistently lower in OsEPF1OE lines than their control, and higher in OsEPFL9OE lines compared to their controls.

|  | Layer 1 | Layer 2 | Layer 3 | Layer 4 | Layer 5 | ANOVA |
| --- | --- | --- | --- | --- | --- | --- |
| **IR64** | 1.22(±0.01)<br>a A | 1.30(±0.01)<br>a B | 1.30(±0.01)<br>a B | 1.28(±0.01)<br>a B | 1.25(±0.01)<br>a A,B | p < 0.0001 |
| **OsEPF1OE-W** | 1.13(±0.01)<br>b A | 1.16(±0.01)<br>b A,B | 1.17(±0.01)<br>b B | 1.14(±0.01)<br>b A,B | 1.14(±0.01)<br>b A,B | p = 0.0140 |
| **OsEPF1OE-S** | 1.14(±0.01)<br>b A | 1.19(±0.01)<br>b B | 1.17(±0.01)<br>b A,B | 1.16(±0.01)<br>b A,B | 1.15(±0.01)<br>b A | p = 0.0060 |
| **ANOVA**<br>**(OsEPF1OE)** | p < 0.0001 | p < 0.0001 | p < 0.0001 | p < 0.0001 | p < 0.0001 |  |
| **IR64** | 1.16(±0.01)<br>a A | 1.20(±0.01)<br>a A,B | 1.22(±0.01)<br>a B | 1.17(±0.01)<br>a A | 1.19(±0.01)<br>a A,B | p = 0.0043 |
| **OsEPFL9OE-2** | 1.25(±0.01)<br>b A | 1.33(±0.01)<br>b B | 1.35(±0.02)<br>b B | 1.27(±0.01)<br>b A | 1.27(±0.01)<br>b A | p < 0.0001 |
| **OsEPFL9OE-3** | 1.25(±0.02)<br>b A | 1.34(±0.02)<br>b C | 1.31(±0.01)<br>b B,C | 1.29(±0.01)<br>b A,B,C | 1.27(±0.01)<br>b A,B | p = 0.0006 |
| **ANOVA**<br>**(OsEPFL9OE)** | p < 0.0001 | p < 0.0001 | p < 0.0001 | p < 0.0001 | p < 0.0001 |  |

**Table 3:**
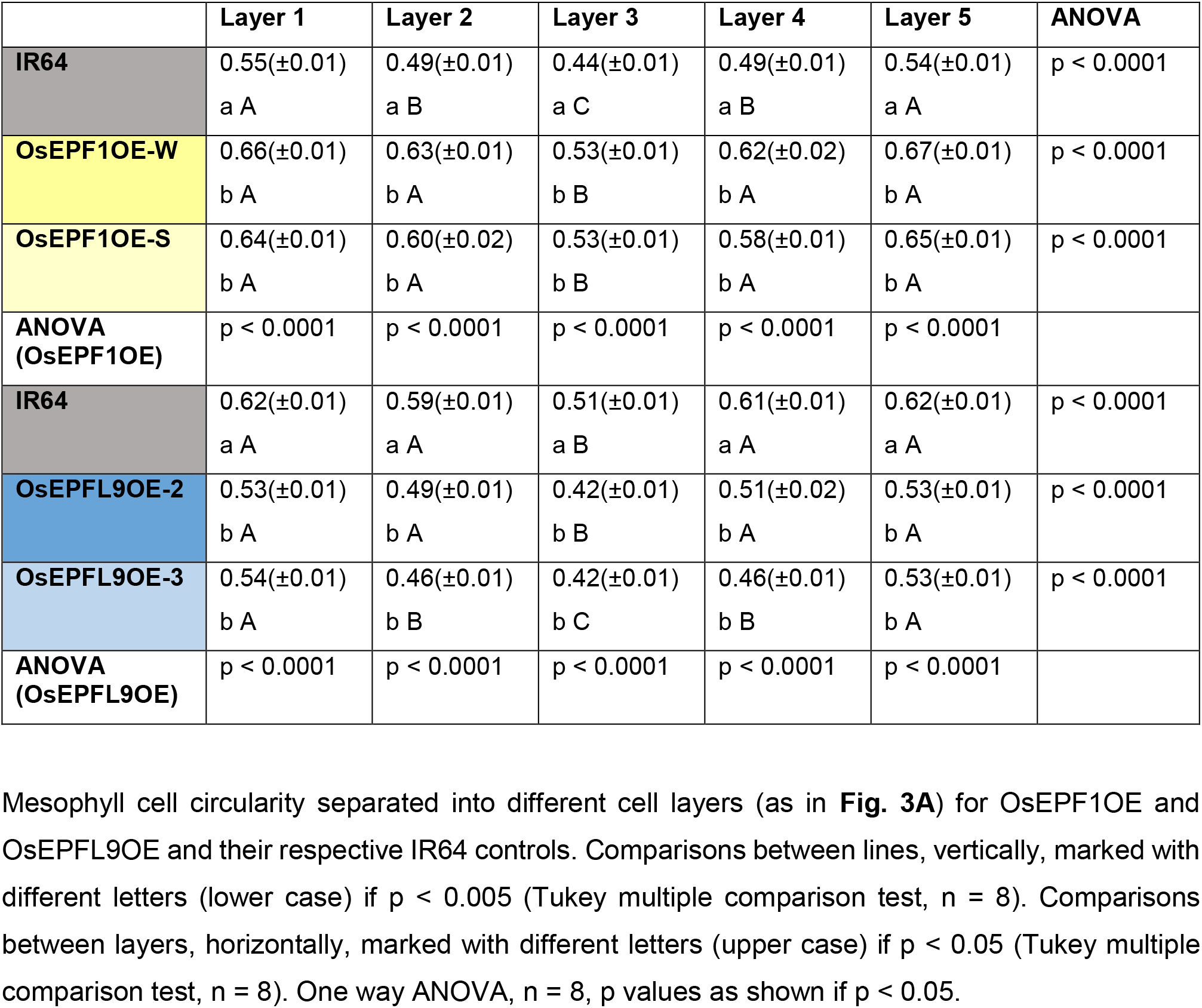
OsEPF1OE and OsEPFL9OE mesophyll cell circularity varies by cell layer, with layer 3 consistently being the least circular. Cell circularity is consistently higher in OsEPF1OE lines than their control, and lower in OsEPFL9OE lines compared to their controls.

To find out whether stomatal density affects the size and shape of all cells throughout the mesophyll in a similar way, we compared the cells in each layer across the six lines (**Tables 1, 2 and 3, Supplementary Fig. S6**). OsEPF1OE cell area was significantly lower than the control in layers 1, 2 and 5 (**Supplementary Fig. S6A**), while cell area in OsEPFL9OE lines was not significantly different from the control in any individual layer (**Table 1, Supplementary Fig. S6B**). Cell lobing was significantly lower in every mesophyll cell layer of OsEPF1OE lines compared to the control, while OsEPFL9OE lines have significantly higher cell lobing than the control in every mesophyll layer (**Table 2, Supplementary Fig. S6C**,**D**, one way ANOVA, all layers: p < 0.0001, Tukey multiple comparisons test p < 0.0001, n = 8). Cell circularity is also affected in the same way in every layer of the mesophyll, with OsEPF1OE mesophyll cells having higher circularity than the control, and OsEPFL9OE mesophyll cells being less circular (**Table 3, Supplementary Fig. S2E**,**F**, one way ANOVA, all layers: p < 0.0001, Tukey multiple comparisons test p < 0.0001, n = 8).

### Patterning of mesophyll cell size and shape is observed in a range of rice cultivars and species

Having established that a pattern of cell size and shape existed in IR64 plants with varying stomatal densities, we were then interested in the universality of this mesophyll patterning in the wider rice family. We therefore studied the mesophyll in a range of rice varieties, including three *O. sativa* Indica variants (MRQ76, MR220 and Malinja), and three wild varieties (*O. latifolia, O. punctata* and *O. meridionalis*). These variants show a range of plant structure and size (**Supplementary Fig. S7**).

Again, one representative variety has been shown in the main text, *O. latifolia*, where the differences between the cell layers are particular clear (**Fig. 4**), with the data from the remaining five varieties shown in the Supplementary Material (**Supplementary Fig. 8-11**) - the same patterns are seen in all varieties. Mesophyll cell size varied by layer in all varieties analysed (one way ANOVA p < 0.05 – p < 0.0001, n = 4-6), and cells were always largest in layer 3 and smallest in layer 4 (**Fig. 4A, Supplementary Fig. S8, Table S1**). The cells in layer 3 were significantly larger than those in all other layers for *O. latifolia* and *O. punctata* (**Fig. 4A, Supplementary Fig. S8D**, Tukey multiple comparison test, p = 0.0002 – p < 0.0001, n = 5, n = 4, respectively). Mesophyll cell lobing (**Fig. 4B, Supplementary Fig. S9, Table S2**) was not significantly different across the adaxial/abaxial axis (one way ANOVA, p > 0.05, n = 4-6), although there was a tendency for lower lobing in the outer layers, particularly layer 1. In *O. latfolia*, mesophyll cell circularity varied significantly by layer (**Fig. 4C**, one way ANOVA, p = 0.0105, n = 8). In all the other rice varieties, circularity was not significantly affected by layer (**Supplementary Fig. S10, Table S3**), but layer 3 did consistently have the lowest circularity. Mesophyll cell projections allow us to see differences in cell shape and size between the tissue layers in all rice varieties (**Fig. 4D, Supplementary Fig. S11**).

### The long axis orientation of mesophyll cells varies with leaf layer position

To investigate whether a pattern in cell alignment also accompanies the layering of mesophyll cell size and shape described above, we analysed the orientation of the mesophyll cell long-axis (see **Supplementary Fig. S12**), in the OsEPF1OE and OsEPFL9OE lines and the IR64 background (**Fig. 3F, Supplementary Fig. 13**) and the range of rice cultivars and species (**Fig. 4F, Supplementary Fig. S14**). An angle of 0° indicates that the longest axis of the cell is horizontal (along the plane of the leaf lamina), whereas cells with an angle of 90° are longest in the vertical plane (perpendicular to the plane of the leaf lamina). The data show that there is a pattern of cell orientation across the adaxial/abaxial axis. The long axis of mesophyll cells in the middle layers (layers 2-4) are noticeably more horizontal than the cells closest to either epidermis (layers 1 and 5). Mesophyll cells in layer 1 do not have a clear dominant cell angle, whereas layer 5 cells appear to have an average cell axiality of between 30 and 45°.

## DISCUSSION

### Linking stomatal function and mesophyll structure in rice

Recent data in Arabidopsis and wheat suggest that the internal structure of the mesophyll is modulated by the activity of stomata on the epidermal surfaces of leaves (Dow *et al*., 2017; Lundgren *et al*., 2019; Wilson *et al*., 2021). The results presented here from rice supports this hypothesis - rice leaves manipulated to have an increased stomatal density and, hence, increased *g*_*s*_ and *gs*_*max*,_ displayed an in increase in cell lobing, with the opposite being true for leaves with decreased stomatal density. In the leaves with reduced cell lobing (OsEPF1OE plants) there was also a decrease in cell size, whereas in the leaves displaying an increased cell lobing (OsEPFL9OE plants) there was no accompanying increase in cell size. This means that in the latter case the change in lobing must reflect a true shape change, whereas in the former case we cannot discount an indirect effect on lobing due to the change in cell size. Changes in mesophyll cell size and/or lobing will influence their surface area/volume ratio, which would likely alter the potential gas flux through the mesophyll. Thus, the increased cell lobing observed in the OsEPFL9OE leaves provides a relative increase in area for gas diffusion into and out of the cell, linking to an expected increase in gas flux due to the increased stomatal density on these leaves. Conversely, the decreased cell lobing observed in the OsEPF1OE leaves provides a relative decrease in area for gas diffusion into and out of the cell, linking to an expected decrease in gas flux due to the decreased stomatal density.

Mesophyll cell lobing in rice has been long associated with the potential for maximising gas flux (Sage and Sage, 2009) and our data support this proposal. However, the mechanism of mesophyll cell lobing and its regulation remains unclear (Lundgren and Fleming, 2020). There is accumulating data on how epidermal cells form intricate lobes to generate the classical jigsaw pattern of this tissue, with the link from cytoskeleton to local wall deformation being established (Sampathkumar *et al*., 2014) and with buckling of the perimeter being postulated as part of the lobe initiation process (Bidhendi and Geitmann, 2019). Presumably, similar molecular processes underpin the control of number and degree of lobing in grass mesophyll cells. Yet, how this process could be modulated by gas flux, and how the surface area/cell volume rheostat is sensed and linked to e.g. photosynthesis, remains to be elucidated.

It is interesting that although the data presented here for rice and previously published for wheat (Wilson *et al*., 2021) both support a role for stomatal-derived gas exchange influencing mesophyll cell size and shape, the fine cellular details (and thus mechanism) of the response may be distinct. In wheat, mesophyll cell volume is larger (with an increase in lobe number) in genotypes with increased stomatal conductance, whereas in rice the overall cell size is little changed but there is a clear change in cell lobing and circularity. Thus, It is possible that different grass leaves employ slightly different cellular approaches to maintaining surface area/volume, a trait which is presumably under evolutionary selection pressure due to its influence on leaf photosynthetic capacity and water loss.

### Rice mesophyll displays a conserved pattern of size and shape

An interesting and unexpected observation resulting from our analysis of mesophyll cell size and shape in mutants with altered stomatal density was the apparent pattern between the geometry of mesophyll cells and their location within the leaf. Quantitative comparison confirmed this was the case, with the middle cells (layer 3) always being larger than cells in adjacent and sub-adjacent layers. Cells in this layer were also characterised as having the lowest degree of circularity and an axiality, which was more parallel to the plane of the leaf surface than cells in the other layers. A distinctive pattern of cell axiality was also observed in the most adaxial mesophyll layer (layer 1) where cells displayed a much wider range than cells in the other layers, and the longest plane of cells in layer 5 was often ∼45°, reflecting their more square shape. Interestingly, these cellular patterns were generally also observed in a range of rice species and cultivars beyond the IR64 lines used for the transgenics, suggesting that the patterns reflect a widespread phenomenon. Moreover, in the transgenic IR64 lines with altered stomatal densities, although the absolute values of some parameters, for example, cell size, shifted (as described in the previous section) the underlying cell patterns remained, suggesting that the stomatal-related signal was modulating an endogenous developmental pattern that was embedded in the leaves.

These observations are in contrast to a text-book view that in monocots mesophyll cell size is distributed uniformly within the leaf (Esau, 1965; Chonan, 1978). There have been previous suggestions that this might be an over-simplification of the true situation. For example, in the original paper highlighting the potential importance of cell lobing (Sage and Sage, 2009), the authors showed that the cells towards the middle of the mesophyll tend to be more elongate, had a larger vacuole, and a lower proportion of chloroplast by volume than cells nearer the outside of the leaf. Our data build and extend this view to show that there is a clear and consistent pattern in a range of rice species in which cells in the middle layer are significantly larger than cells in other layers of the mesophyll, have a distinct shape (higher circularity) and display a restraint in cell axiality absent in cells in other layers of the leaf. Borsuk et al. (2022) recently used microCT technology to show that the dicotyledonous spongy mesophyll is also more organised than was previously thought, suggesting that more modern techniques and thorough analysis may be discovering patterns in leaf tissues which were previously considered disordered.

These observations lead to the question of how the rice mesophyll pattern arises and what, if any, advantage this arrangement of cells conveys to the leaf. With respect to development, Zeng et al. (2016) showed that the middle layer of the rice mesophyll (layer 3 in this paper) is derived from the L3 cells of the leaf primordium, whereas the cells neighbouring the epidermal cells are derived from L2 cells. The layer 3 cells are thus likely to be clonally distinct, so that their size, shape and axiality might, theoretically, reflect their ontogeny. A more precise analysis of cell size and shape across the emerging layers in the developing rice leaf would help test this possibility. An alternative (though not exclusive) hypothesis is that the cellular pattern across the adaxial/abaxial axis of the leaf is linked to specific function. For example, in many eudicot leaves the mesophyll cells that form the distinct palisade layer are vertically aligned and cylindrical in shape to aid light penetration to the lower spongy mesophyll (Vogelmann *et al*., 1996). It is possible that the more vertical orientation of the cells in layer 1 and 5 of the rice mesophyll (the external layers of the mesophyll) have a similar role in directing light towards the more internal mesophyll of the leaf. Investigating light distribution in leaves with a range of layer 1 cell axiality might help distinguish these possibilities. In a similar way, the horizontally elongate layer 3 cells could be specialised to, for example, transport solutes between veins.

Another possibility is that the variation in cell size and shape across the mesophyll reflects a trade-off between optimising surface/area to volume for gas exchange, the optimum spread of material for light absorption, and the investment costs (carbon, nitrogen, energy) in building a leaf, as has been explored by (Earles *et al*., 2019). In order to investigate this idea we have created four simplified models of mesophyll cell packing (**Fig. 5A-D**). Two different cell types were used based on the length and width measurements from *O. latifolia* (**Supplementary Fig. S15**). Cell wall thickness and mitochondria size and distribution are the same for both cell types, but large cells have a lower proportion of plastid and higher proportion of cytosol, to reflect the findings of Sage and Sage (2009). Model 1 is most representative of mesophyll described in this study, with larger cells in the middle layer (layer 3) (**Fig. 5A**). Model 2 has five layers of small cells, making the ‘leaf’ slightly thinner than Model 1 (**Fig. 5B**). Model 3 has the same ‘leaf’ thickness as Model 2, but is made up of four layers of large cells (**Fig. 5C**). Model 4 is also made entirely or large cells, but has five layers of cells resulting in the same plastid and cytosol volume as Models 1 and 2 (**Fig. 5D**). Models 2 and 3 have the same ‘leaf’ thickness, while Model 4 is the thickest. One layer of three large cells has the same plastid and cytosol volume as one layer of five small cells. The amount of cell wall in contact with the air (*S*_*mes*_) is very similar in Models 1 and 2, lowest in Model 3 and intermediate in Model 4 (**Fig. 5E**). When the model leaves are supplied with incident light from the adaxial surface, Models 1 and 2 have higher total light absorptance than Models 3 and 4 (**Fig. 5F**). However, Models 3 and 4 (consisting of entirely larger cells) allow more light to travel further into the leaf, with significantly higher absorptance than Model 1 in cell layers 3 and 4 (**Fig. 5G**). This can be explained by the stronger sieve effect (as in Terashima et al., 2009) in the large cells due to the chloroplast being spread more sparsely. Modelled photosynthetic performance was similar between the four cell tissue layer models, although Models 3 and 4 do perform slightly less well, particularly during the Rubisco-limited initial slope of the curve (**Fig. 5H**). Unsurprisingly, Model 3, with the lowest volume of chloroplast and smallest light absorptance has the lowest assimilation at low internal CO_2_. Our models suggest that, with respect to light absorption and photosynthesis there is little to distinguish Model 1 and Model 2 (with the proviso, of course that these models represent major simplifications of the system). Allowing for these limitations, if light absorption and photosynthesis are not the functional drivers for the pattern of larger, more horizontally aligned cells in layer 3, what might the function be? At present we can only speculate. For example, it might reflect a mechanical role in supporting the leaf lamina. Alternatively, a by-product of the pattern is fewer cell boundaries in the lateral plane of the leaf connecting adjacent veins. If layer 3 has a role in transporting molecules to and from vascular bundles, a trait of fewer cell boundaries might be advantageous.

Finally, our findings have implications (both negative and positive) for related research in the broader area of rice research. Firstly, many studies taking a comparative approach to leaf structure in grasses use the middle layer of the mesophyll as an easily identifiable region to sample, thus decreasing the work-load involved in often largescale analyses (e.g. Ouk *et al*., 2020). Our data suggest that, unfortunately, the cells in this layer are in some ways atypical of the mesophyll as a whole. On the other hand, there is significant interest in engineering rice leaves to instil a major shift in photosynthesis (C_4_ photosynthesis) - with decreasing the number of mesophyll cells between vascular bundles as a key aim (Ermakova *et al*., 2020). Our data indicate that, due to their size and axiality, the middle layer of the rice mesophyll already provides the fewest cells between neighbouring veins. Driving this anisotropic growth further is an avenue to explore which might contribute to achieving this leaf engineering goal.

## ABBREVIATIONS

EPF(L): Epidermal Pattering Factor (Like)
*g*_*s*_: stomatal conductance
*g*_*smax*_: maximum theoretical stomatal conductance
*S*_*mes*_: surface area of mesophyll cell in contact with air

## SUPPLEMENTARY DATA

Supplementary data are available at *JXB* online.

Figure S1: Stomatal density and theoretical *g*_*smax*_ is altered in EPF1 and EPFL9 OE lines

Figure S2: Layer 3 mesophyll cells are larger than other cell layers

Figure S3: Layer 1 mesophyll cells have the lowest values of lobing

Figure S4: Layer 3 mesophyll cells have the lowest circularity

Figure S5: Mesophyll cell projections show the variety of cell shapes and sizes in the mesophyll cell tissue layers in EPF1OE, EPFL9OE and IR64 control lines

Figure S6: Mesophyll cell area, lobing, circularity by layer

Figure S7: Six different varieties of rice used in Figure 4 and Figures S8-11 show a range of plant structure and size

Figure S8: Layer 3 mesophyll cells are the largest across a range of rice varieties

Figure S9: Layer 1 mesophyll cells always have the lowest lobing value across a range of varieties

Figure S10: Layer 3 mesophyll cells have the lowest circularity across a range of rice varieties

Figure S11: Mesophyll cell projections show the variety of cell shapes and sizes in the mesophyll cell tissue layers in a range of rice varieties

Figure S12: Measurement of mesophyll cell lobing and orientation

Figure S13: Internal layers of mesophyll cells have a more horizontal long axis in EPF1OE, EPFL9OE and IR64 control lines

Figure S14: Internal layers of mesophyll cells have a more horizontal long axis in a range of six rice varieties

Figure S15: Measurements of large and small cells used in leaf tissue models

Table S1: Mesophyll cell area in a range of six rice varieties

Table S2: Mesophyll cell lobing in a range of six rice varieties

Table S3: Mesophyll circularity in a range of six rice varieties

## ACKNOWLEDGEMENTS

J.S. and Y.X. were supported by a Royal Society Challenge-Led grant CHL\R1\18007 “Breeding rice resilient to a high CO_2_ future” to A.J..F and X.-G.Z.. J.A. was supported by the BBSRC White Rose DTP (BB/T007222/1). S. I.-C. was supported by a PhD studentship from the Thai government. M.J.W. was supported by a BBSRC grant Shape Shifting Stomata: The Role of Geometry in Plant Cell Function (BB/T005041) to A.F.. Thank you to Rachel Thorley for preliminary discussions on cell tissue layers.

## AUTHOR CONTRIBUTIONS

J.S., S.I.-C., Q.Y.N., J.A. and M.J.W. performed the experiments; Y.X. performed the computational modelling, J.S., S.I.-C., Y.Q.N., Y.X., J.A., M.J.W., X.-G.Z. and A.J.F interpreted the results and wrote the paper, with contributions from all authors. A.J.F. designed the study and led the project.

## DATA AVAILABILITY

The data supporting the findings of this study are available from the corresponding author, (Dr Jen Sloan), upon request.

## FIGURE LEGENDS

**Figure S1: stomatal density and theoretical *g***_***smax***_ **is altered in EPF1 and EPFL9 OE lines**

Data from the middle of leaf 6. **A**,**C**,**E)** 28 day old OsEPF1OE weak (W) and strong (S) lines, and their IR64 control. **B**,**D**,**F)** 21 day old OsEPFL9OE line 2 and 3 and their IR64 control. **A)** OsEPF1OE abaxial stomatal density is significantly lower than in the control. One way ANOVA, p < 0.0001, n = 8. **B)** OsEPFL9OE abaxial stomatal density is significantly higher than in the control. One way ANOVA, p = 0.0002, n = 8. **C)** OsEPF1OE guard cell length is not different from the control. One way ANOVA, p > 0.05, n = 8. **D)** OsEPFL9OE-3 has significantly smaller guard cells than the control. One way ANOVA, p = 0.0051, n = 8. **E)** OsEPF1OE lines have significantly lower theoretical *g*_*smax*_ than the control. One way ANOVA, p < 0.0001, n = 8. **F)** OsEPFL9 lines have significantly higher theoretical *g*_*smax*_ than the control. One way ANOVA, p = 0.0028.

All multiple pairwise comparisons, Tukey, p values as shown, n= 8.

**Figure S2: Layer 3 mesophyll cells are larger than other cell layers**

Mesophyll cell area from the middle of leaf 6. **A)** 28 day old EPF1 IR64 control, **C)** OsEPF1OE-W and **E)** OsEPF1OE-S, **B)** 21 day old EPFL9 IR64 control, **D)** OsEPFL9OE-2 and **F)** OsEPFL9OE-3. Cell area varies in the adaxial/abaxial plane, one way ANOVA, p < 0.0001 or p = 0.0001 (see Table 1), n = 8. Layer 3 cells are significantly larger than cells in the other layers in all lines, multiple pairwise comparisons, Tukey, p values as shown, n = 8.

**Figure S3: Layer 1 mesophyll cells have the lowest values of lobing**

Mesophyll cell lobing from the middle of leaf 6. **A)** 28 day old EPF1 IR64 control, **C)** OsEPF1OE-W and **E)** OsEPF1OE-S, **B)** 21 day old EPFL9 IR64 control, **D)** OsEPFL9OE-2 and **F)** OsEPFL9OE-3. Cell lobing varies in the adaxial/abaxial plane, one way ANOVA, p < 0.0001-p = 0.014 (see Table 2), n = 8. Layer 1 cells always have the lowest lobing level, multiple pairwise comparisons, Tukey, p values as shown, n = 8.

**Figure S4: Layer 3 mesophyll cells have the lowest circularity**

Mesophyll cell circularity from the middle of leaf 6. **A)** 28 day old EPF1 IR64 control, **C)** OsEPF1OE-W and **E)** OsEPF1OE-S, **B)** 21 day old EPFL9 IR64 control, **D)** OsEPFL9OE-2 and **F)** OsEPFL9OE-3. Cell circularity varies in the adaxial/abaxial plane, one way ANOVA, p < 0.0001, n = 8. Layer 3 cells have significantly lower circularity than cells in the other layers in all lines, multiple pairwise comparisons, Tukey, p values as shown, n = 8.

**Figure S5: Mesophyll cell projections show the variety of cell shapes and sizes in the mesophyll cell tissue layers in EPF1OE, EPFL9OE and IR64 control lines**

Mesophyll cell projections of all cells in each layer from one representative individual per geneotype. Scale bar = 20 μm

**Figure S6: Mesophyll cell area, lobing, circularity by layer**

**A**,**C**,**E)** 28 day old leaf 6 from OsEPF1OE lines and their control. **B**,**D**,**F)** 21 day old leaf 6 from OsEPFL9OE lines and their control. **A**,**B)** Mesophyll cell area by layer. **C**,**D)** Mesophyll cell lobing by layer, **E**,**F)** Mesophyll cell circularity by layer.

**A)** One way ANOVA, layer 1: p = 0.0073, layer 2: p = 0.0003, layer 3: p = 0.0340, layer 4: ns, layer 5: p = 0.0015, n = 8. **B)** One way ANOVA, layer 1: ns, layer 2: p = 0.0012, layer 3: ns, layer 4: p = 0.0207, layer 5: ns, n = 8. **C)** One way ANOVA, all layers: p < 0.0001, n = 8. **D)** One way ANOVA, all layers: p < 0.0001, n = 8. **E)** One way ANOVA, all layers: p < 0.0001, n = 8. **F)** One way ANOVA, all layers: p < 0.0001, n = 8.

**Figure S7: Six different varieties of rice used in Figure 4 and Supplementary Figures 8-11 show a range of plant structure and size**

Plants pictured at 35 days old. **A)** *O. sativa* (MR220), **B)** *O. latifolia* **C)** *O. sativa* (MRQ76), **D)** *O. punctata*, **E)** *O. sativa* (Malinja) **F)** *O. meridionalis*

**Figure S8: Layer 3 mesophyll cells are the largest across a range of rice varieties**

Mesophyll cell area from the middle of leaf 6 of six rice varieties. Cell size varies across the adaxial/abaxial axis in all varieties. One way ANOVA: **A)** *O. sativa* (MR220), p = 0.0081 n = 6, **B)** *O. latifolia*, p < 0.0001, n = 6, **C)** *O. sativa* (MRQ76), p = 0.0368, n = 5, **D)** *O. punctata*, p < 0.0001, n = 4, **E)** *O. sativa* (Malinja), p = 0.0009, n = 6, **F)** *O. meridionalis*, p = 0.0467, n = 6. Cells in layer 3 are largest and layer 4 cells are smallest in every variety. In *O. latifolia* **(B)** and *O. punctata* **(D)**, layer 3 cells are significantly larger than cells in any other layer.

All multiple pairwise comparisons, Tukey, p values as shown, n = 4-6.

**Figure S9: Layer 1 mesophyll cells always have the lowest lobing value across a range of varieties**

Mesophyll cell lobing from the middle of leaf 6 of six rice varieties – **A)** *O. sativa* (MR220), **B)** *O. latifolia* **C)** *O. sativa* (MRQ76), **D)** *O. punctata*, **E)** *O. sativa* (Malinja) **F)** *O. meridionalis* Cell lobing does not significantly vary across the abaxial/adaxial gradient. One way ANOVA, p > 0.05, n = 4-6. Cells in layer 1 always show the lowest level of lobing.

**Figure S10: Layer 3 mesophyll cells have the lowest circularity across a range of rice varieties**

Mesophyll cell area from the middle of leaf 6 of six rice varieties – **A)** *O. sativa* (MR220), **B)** *O. latifolia*, One way ANOVA, p = 0.0105, Tukey multiple pairwise comparisons, p values as shown, n = 6, **C)** *O. sativa* (MRQ76), **D)** *O. punctata*, **E)** *O. sativa* (Malinja) **F)** *O. meridionalis*. **A**,**C**,**D**,**E**,**F)** One way ANOVA, p > 0.05, n = 4-6

**Figure S11: Mesophyll cell projections show the variety of cell shapes and sizes in the mesophyll cell tissue layers in a range of rice varieties**

Mesophyll cell projections of all cells in each layer of one representative individual for 5 rice varieties Scale bar = 20μm

**Figure S12: Measurement of mesophyll cell lobiness and orientation**

**A)** a line was drawn between the two minor veins in each image. The angle of this line was measured and considered horizontal.

**B)** Cell perimeter and convex hull perimeter were measured in ImageJ. Lobiness is calculated as cell perimeter/convex hull perimeter.

The FeretAngle measurement (0-180 degrees) is the angle between the Feret’s diameter and a line parallel to the x-axis of the image. The horizontal angle was subtracted from this angle so that a cell angle of 0° is parallel to the line between the minor veins. If the FeretAngle is >180°, the angle was adjusted (180-FeretAngle) so that all angles were between 0 and 90° for ease of comparison. A cell with an angle of 90°is aligned with its longest axis vertical (or perpendicular to the line between the minor veins).

**Figure S13: Internal layers of mesophyll cells have a more horizontal long axis**

Mesophyll cell angle in cells from different cell layers from the middle of leaf 6. 28 day old EPF1 IR64 control, OsEPF1OE-W and OsEPF1OE-S, 21 day old EPFL9 IR64 control, OsEPFL9OE-2 and OsEPFL9OE-3. The longest axis of cells in the internal mesophyll layers (2-4) is more horizontal than the layers adjacent to the epidermes. Cells in layer 1 (adaxial) have a fairly random distribution of cell angle, layer 5 cells (abaxial) are most commonly at an angle of 30-40°.

**Figure S14: Internal layers of mesophyll cells have a more horizontal long axis in a range of six rice varieties**

The longest axis of cells in the internal mesophyll layers (2-4) is more horizontal than the layers adjacent to the epidermes. Cells in layer 1 (adaxial) have a fairly random distribution of cell angle, layer 5 cells (abaxial) are most commonly at an angle of 30-40°.

**Figure S15: Measurements of large and small cells used in leaf tissue models**

**A)** Detailed representation of each cell in the leaf tissue model. **B)** Different parameter measurements used for small and large cells in leaf tissue models

**Table S1: Mesophyll cell area in a range of six rice varieties**

Mesophyll cell area separated into different cell layers (as in **Fig. 3A**) for six *Oryza* varieties. Comparisons between lines, vertically are not significant (one way ANOVA, p > 0.05, n = 4-6). Comparisons between layers, horizontally, marked with different letters (upper case) if p < 0.05 (Tukey multiple comparison test, n = 4-6). One way ANOVA, n = 4-6, p values as shown.

**Table S2: Mesophyll cell lobiness in a range of six rice varieties**

Mesophyll cell lobiness separated into different cell layers (as in **Fig. 3A**) for six *ORYZA* varieties. Comparisons between lines (vertically), and between layers (horizontally) were not significant (One way ANOVA, p > 0.05, n = 4-6.

**Table S3: Mesophyll circularity in a range of six rice varieties**

Mesophyll cell circularity separated into different cell layers (as in **Fig. 3A**) for six *ORYZA* varieties. Comparisons between lines (vertically), and between layers (horizontally) were not significant (One way ANOVA, p > 0.05, n = 4-6.

